# Deformation Gradient Tensor Model of Roll–Spiral Transformation for Protein Assembly R-Body

**DOI:** 10.64898/2026.05.26.728034

**Authors:** Satoru Tsugawa, Kosuke Kikuchi, Koki Date, Tomonobu Nonoyama, Zichen Kang, Takafumi Ueno

## Abstract

Spiral geometries commonly occur in natural and engineered systems and are fundamentally described by curvature and torsion. In deformation-dominated systems, these variables evolve dynamically, requiring a continuum mechanical framework to link geometry and deformation. This study focused on refractile bodies (R-bodies), protein supramolecular assemblies that undergo reversible roll–spiral transformations in response to stimuli such as pH changes. Although multiple R-body types with distinct morphologies and unrolling behaviours were experimentally identified, their deformation mechanisms lack quantitative theoretical descriptions. We proposed a deformation-gradient-tensor-based continuum model incorporating geometrical mapping from the rolled to spiral state within a unified framework. The model successfully reconstructed macroscopic deformation behaviours of types 51, 7, and Pa R-bodies by capturing differences in unrolling behaviours, tapered geometry, and spatio-temporal evolution. The analysis revealed that deformation proceeds through a coupled process in which the curvature decreases via straightening, while the torsion increases by twisting. Importantly, the framework connected the macroscopic morphology with microscopic lattice deformation, enabling quantitative inference of lattice intervals and angles. The proposed comprehensive geometric model of the R-body roll–spiral transformation offers a general mathematical foundation for understanding deformation-driven spiral transformations in soft matter systems.

## 1. Introduction

Spiral geometries are ubiquitous throughout nature and engineering, spanning the molecular, cellular, and macroscopic scales. Such structures appear in diverse systems including bacterial spirochetes (Goldstein 1988), seashells (Moulton 2012, Moulton 2014), plant tendrils (Sachs 1875), engineered helical cables (Hearle 1980) and architectural forms (Israel 2015). Despite this diversity, their geometries can ultimately be characterised in a unified manner using two key variables: curvature and torsion (Goriely 2017). Curvature describes the bending within a local tangent plane, whereas torsion describes the twisting relative to the tangent plane (Audoly 2010). In growth-dominated systems such as seashells or antelope horns (Moulton 2012, Moulton 2014, Goriely 2017), curvature and torsion are typically prescribed during formation via anisotropic growth and remain fixed thereafter. In contrast, in deformation-dominated systems such as helical springs or responsive sheets (Iamsaard 2014), these variables evolve dynamically over time owing to three-dimensional deformation. Understanding such dynamic evolution requires a continuum framework that captures both geometry and deformation.

Continuum mechanics offers a mathematical framework for describing spiral geometries arising from growth and deformation (Kaczmarski 2022). In particular, the deformation gradient tensor (DGT) provides a unified description of the positioning and deformation of material elements in space, thereby linking local deformation to the global geometry (Goriely 2017). In deformation-dominated systems, structural changes result from coupled three-dimensional deformation along the axial, circumferential, and thickness directions (Audoly 2010). DGT-based formulations have been successfully applied to a wide range of systems, including the inflation-extension of a tube (Horny 2012) and a half-plane in compression (Goriely 2017). However, despite advances in continuum mechanics, comparable geometric models for bio-macromolecular deformation remain largely absent, except for a few systems, such as bacteriophage T4 (Fraser 2021).

Refractile bodies (R-bodies) are a family of protein supramolecular assemblies that undergo drastic and reversible roll–spiral transformations in response to external stimuli (Preer 1966, Meenaghan 1984, Polka 2016). The type 51 R-body adopts a tightly rolled morphology at neutral pH, whereas acidic conditions induce extension into a long spiral morphology (figure 1*a*–*c*). This deformation generates mechanical forces capable of rupturing cellular membranes, leading to the death of the host cells (Preer 1953, Preer 1966). Since their discovery as intracellular kappa particles in the killer strains of *Paramecium* more than 70 years ago (Sonneborn 1938, Preer 1953), scientists have investigated the molecular basis of R-bodies (Lalucat 1988, Heruth 1994, Date 2026). Several types of R-bodies have been classified on the basis of their macroscopic shape and unrolling direction, such as type 51 R-bodies, which unroll from the inside, and type 7 and Pa R-bodies, which unroll from the outside with blunt or tapered sheet morphologies (figure 1*d*; Preer 1972, Pond 1989). However, in contrast to experimental efforts, a theoretical understanding of R-body deformation has yet to be reported. To date, a quantitative geometric framework has not been proposed for the R-body roll–spiral transformation.

**Figure 1.**
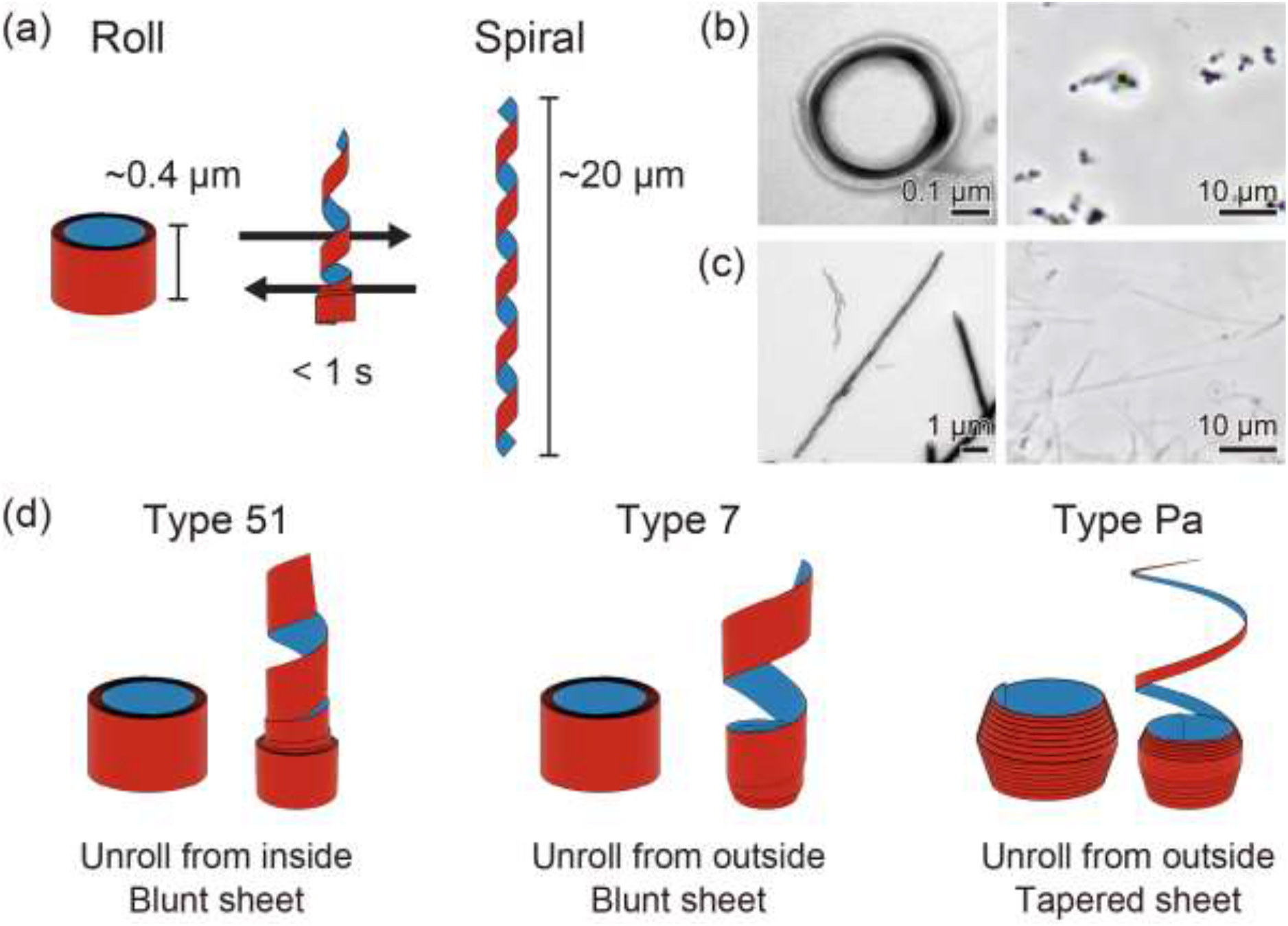
Protein supramolecular ribbon R-body undergoes micrometre-scale, reversible dynamic deformations between the rolled and spiral morphologies. (a) Schematics of type 51 R-body deformations. The outer surface and inner surface are shown in red and blue, respectively. (b-c) Electron micrograph (left) and phase contrast optical image (right) of rolled morphology at pH 7 (b) and spiral morphology at pH 5 (c). (d) Schematics of various shapes and unroll modes of the reported R-body. The type 51 R-body has a blunt sheet and unrolls from the inside (left). The type 7 R-body also has a blunt sheet, but it unrolls from the outside (middle). The type Pa R-body has a tapered sheet toward the edge, of which the outer end is apex-shaped, and unrolls from the outside (right).

In this study, we introduced a DGT-based continuum model for the roll–spiral transformation in an R-body. By formulating a geometrical mapping from roll to spiral within the DGT framework, we describe multiple deformation modes in a unified manner. Using a top-down approach, we modelled macroscopic roll–spiral transformations through the temporal evolution of curvature and torsion. In parallel, a bottom-up analysis relates microscopic lattice deformation to the observed macroscopic geometry. This framework provides the first quantitative geometric description of R-body deformation. More generally, it offers a mathematical foundation for understanding deformation-driven roll–spiral transformations in soft matter systems.

## 2. Results

### 2.1. Development of a DGT model for R-body morphological changes

To quantitatively clarify the morphological descriptions of the R-bodies, we adopted universal deformation models for isotropic materials (Goriely 2017). The rolled and spiral shapes of the R-body can be prescribed by the height, thickness, curvilinear length along the sheet, and the inner and outer diameters of the sheet (figure 2*a*). We set the coordinate systems of the rolled and spiral states as pre-coordinate ***X*** = (*R, Θ, Z*) and post-coordinate ***x*** = (*r, θ, z*) of the centreline of the R-body, respectively (figure 2*b*). A general geometrical mapping between the roll and spiral configurations is expressed as follows:

**Figure 2.**
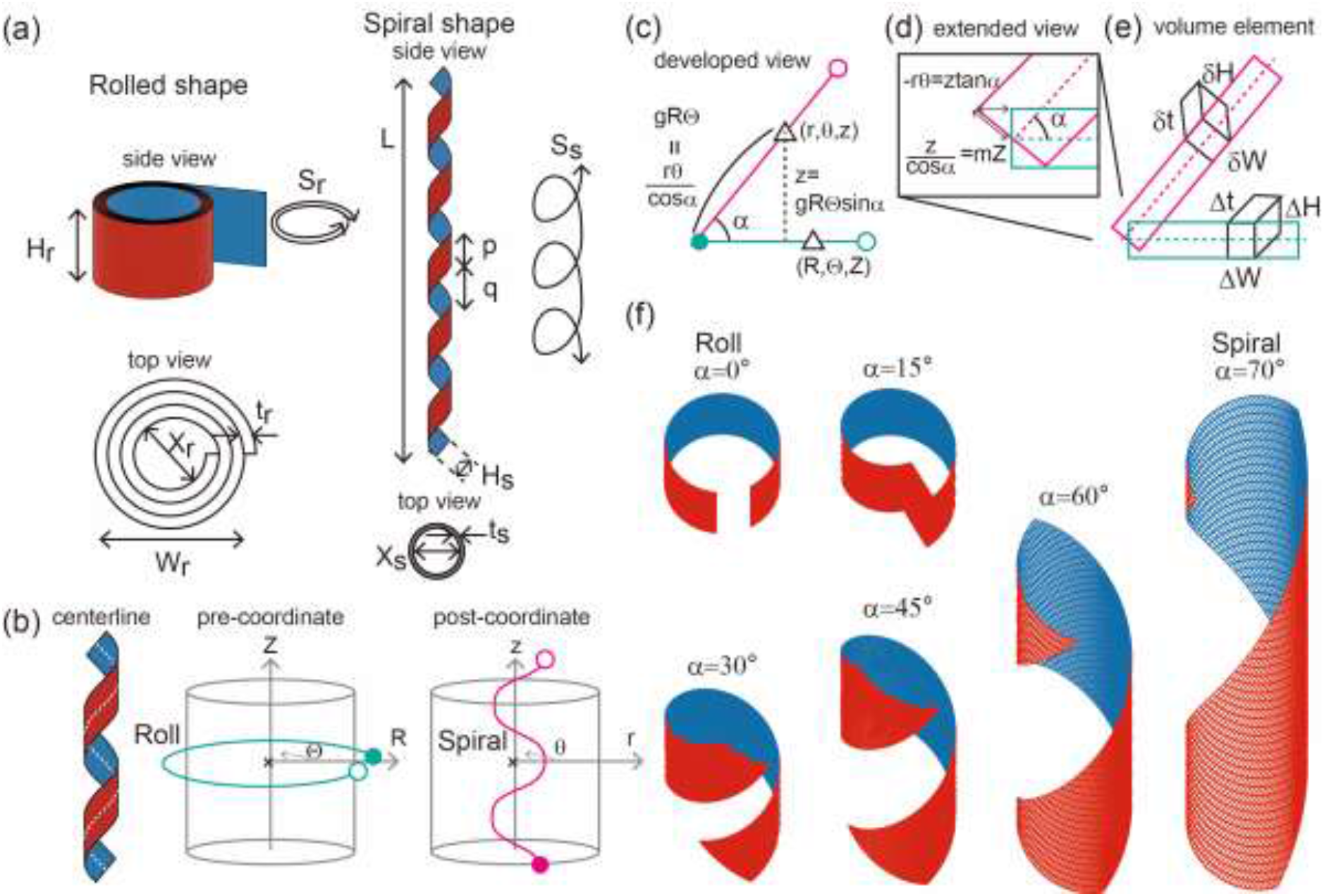
DGT model for the roll–spiral transformation. (a) The rolled shape is determined by height *H*_*r*_, width *W*_*r*_, curvilinear length along the sheet *S*_*r*_, inner diameter *X*_*r*_, and thickness of the sheet *t*_*r*_. The spiral shape is determined by height *H*_*s*_, pitch *p*, interval of pitch *q*, axial length *L*, curvilinear length along the sheet *S*_*s*_, inner diameter *X*_*s*_, and thickness of the sheet *t*_*s*_. (b) Definition of the cylindrical coordinates. From the centreline of the sheet (white dotted curved line on the left), the pre-coordinate system in the rolled state is defined as (*R, Θ, Z*) (centre), and the post-coordinate system at the spiral state is defined as (*r, θ, z*) (right). (c) Developed view of the centreline demonstrating a geometrical relationship. (d, e) Schematic of the volume element around the centreline and (e) extended view of the edge of the sheet (d). (f) Simulated result of the model with different inclination angles *α*. We used *k* = *g* = *m* = 1 and *a* = 1, *b* = 0 in this simulation.

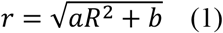

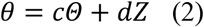

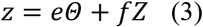

where *a* - *f* are the model parameters. When all the parameters are determined, the deformation between the roll and spiral configurations is completely described.

Using the pre- and post-coordinates, the centreline in the developed view indicates the following two geometrical relationships:

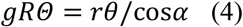

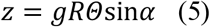

where *g* is the growth parameter in longitudinal length and *α* is the inclination angle of the spiral shape in the developed view. *RΘ* is the arc length of the roll in the pre-coordinate, such that *gRΘ* means the length after deformation, which geometrically corresponds to *rθ*/cos*α* (figure 2*c*). The geometrical relationship near the edge can be written as follows:

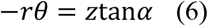

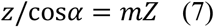

where *m* denotes the height growth parameter in the extended view and −*rθ* is the arc length of the spiral with sign (figure 2*d*). Considering the volume element of the roll and spiral around the centreline as (Δ*H*, Δ*L*, Δ*T*) and (Δ*h*, Δ*l*, Δ*t*), respectively (figure 2*e*), the volumetric change of the element can be calculated as follows:

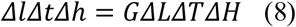

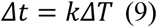

where *G* is the volumetric growth parameter and *k* is the thickness of the growth parameter.

From eqs. (4)–(7), parameters *c* - *f* were determined as follows:

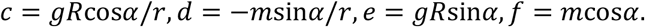

Therefore, the DGT is defined as ***F***= d***x***/d***X***, which can be expressed in the matrix form using eqs (8) and (9),

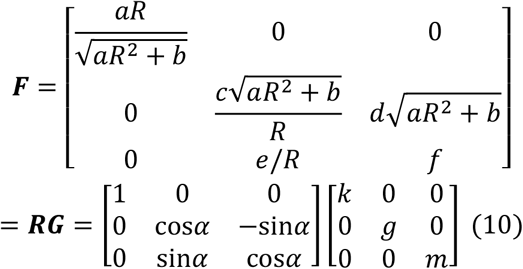

consistent with the work of Goriely (2017). Eq. (10) demonstrates that the roll–spiral transformation of the R-body can be prescribed by two independent matrices: rotation matrix ***R*** around the radial axis by angle *α* and growth matrix ***G*** in three dimensions in the thickness, longitudinal length, and height directions controlled by *k, g*, and *m*, respectively.

### 2.2. Spatio-temporal characterisation of types 51, 7, and Pa R-bodies

Using the DGT model, we built the corresponding parameter sets for the types 51, 7, and Pa R-bodies. To visualise the morphological changes, we discretised the initially twice rolled surface points (*R, Θ, Z*) into 100 and 40 grid points along the circumferential direction and height, respectively. Additionally, we introduced 10 grid points along the thickness direction to represent a finite sheet. We transformed these surface points by applying the DGT model to obtain the deformed coordinates (*r, θ, z*). We note that the surface that had initially been twice rolled can be transformed to an approximate spiral shape that had been rolled once around the principal axis because of the range of variables *Θ* ∈ [0, 4π] rad and *Z* ∈ [0, 200] nm with *c* = 0.68, *d* = −0.01, leading to *θ* ∈ [0, 2π + 0.26] according to eq. (2). In addition, parameter *e* is crucially important for the roll–spiral transformation because parameter values *e* = 188, *f* = 0.34, indicate that parameter *e* has an exceptionally larger value than *c, d*, and *f*. This represents that the system effectively transforms the initial circumferential coordinate *Θ* to the deformed height coordinate *z*.

As a result, types 51, 7, and Pa were reconstructed in a manner consistent with the observed morphology (figure 3*a,b,c*; Gibson 1987, Pond 1989). Type 51 could be reconstructed by prescribing an initial rolled morphology with a gradually decreasing radius. In contrast, types 7 and Pa can be reconstructed by prescribing an initially rolled morphology with a gradually increasing radius and tapered height in type Pa. These changes resulted in opposite behaviours between the inner and outer surfaces. Once inclination angle *α* reached 70°, we discontinued changes to *α* in these simulations, which we used to estimate the experimental images in Pond 1989. We also assumed a time-delayed propagation of the deformation from the starting end to the remainder of the sheet. The type 51 model exhibited an initial deformation of the inner surface. We noticed that the sheet of type 51 gradually unrolled from the inner end, where the striped structure in Cartesian coordinate *x* first disappeared at the inner end (figure 3*d*), and the inner end subsequently turned upward (figure 3*f*). By contrast, the type 7 and Pa models showed that the outer surface starts to deform. Similar to Type 51, the striped structure in *x* disappeared at the outer end (figure 3*e*), and the outer end turned upward (figure 3*g*). Type Pa is almost the same as type 7, except for the tapered height (figure 3*h,i*).

**Figure 3.**
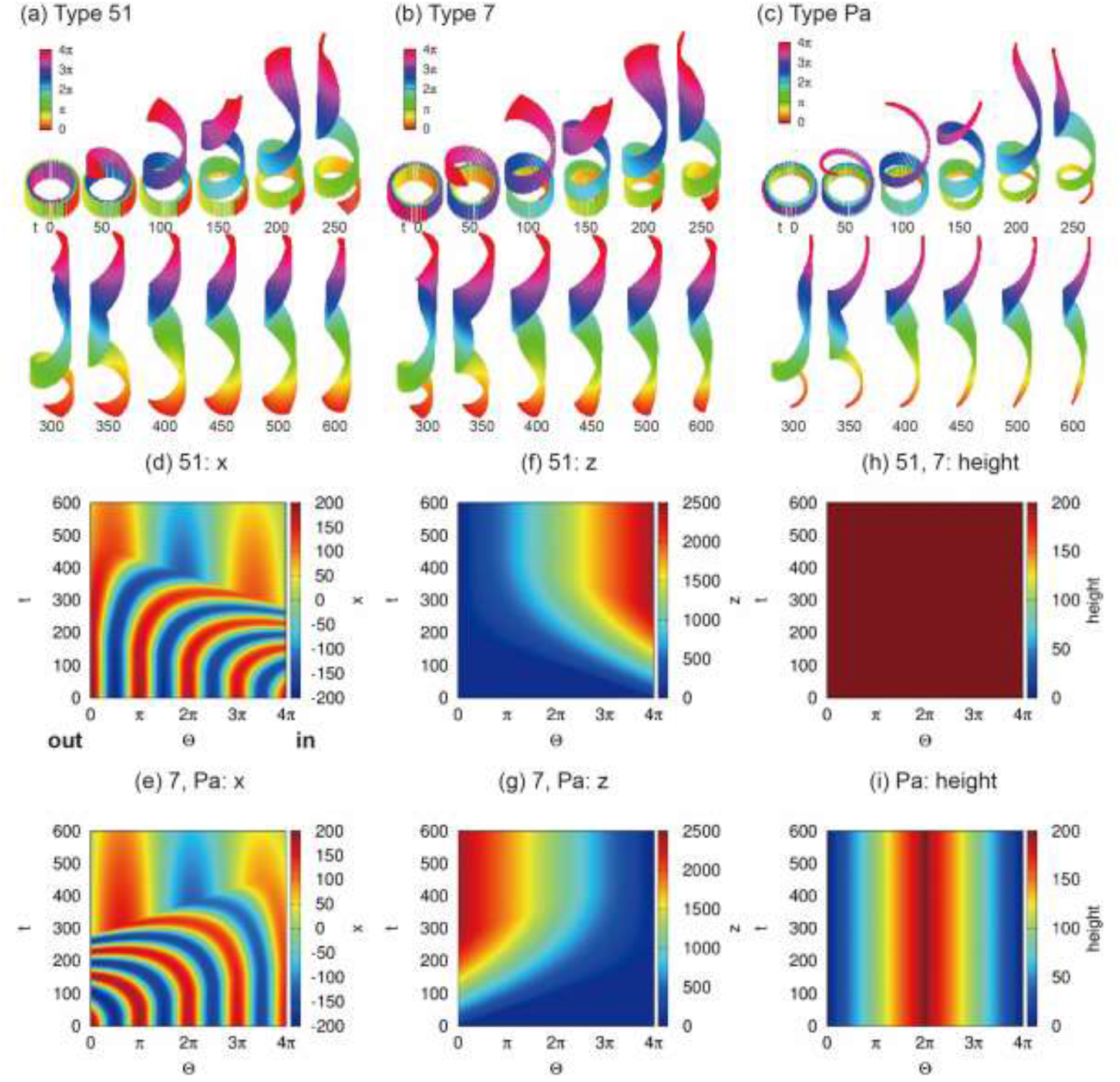
Use of the DGT model to assess different morphological changes in types 51, 7, and Pa. (a, b, c) Simulation results of the type 51 model (a), type 7 model (b), and type Pa model (c). The colour diagram labels initially coordinate *Θ* from 0 to 4π. (d, e) Coordinate *x* as a function of initially labelled coordinate *Θ* along the longitudinal direction and time t of type 51 (d) and types 7 and Pa (e). (f, g) Coordinate *z* of type 51 (f) and types 7 and Pa (g). (h, i) Height of types 51 and 7 (h) and type Pa (i). We used *k* = *g* = *m* = 1 and *α* = 70° (deg) according to the reported image of the R-body (Pond 1989). The initial rolled shape is set with *R* ∈ [150, 200] nm, *Θ* ∈ [0, 4π] rad, *Z* ∈ [0, 200] nm. We assumed *r* ∈ [50,100] nm and set parameters *a* = −1/1200(t − 1200) and *b* = −50*t*/3.

We also calculated the spatio-temporal curvature of the centreline of the sheet, and showed that the curvature of type 51 decreased from the inner end, whereas those of types 7 and Pa decreased from the outer end (figure 4*a,c,b,d*). These results indicated that the maximum bending degree at the dynamically deforming end began to decrease, as inferred from the decreasing curvature. In contrast, the spatio-temporal torsion of type 51 increased from the inner end, and those of type 7 and Pa increased from the outer end, thus converging to a stable torsion (figure 4*e,g,f,h*). These results indicate that the degree of twisting at the dynamically deforming end began to increase, as inferred from the increase in torsion. The obtained results are consistent with macroscopic curvature 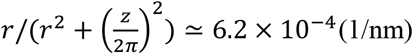 and torsion 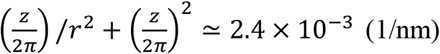 of the spiral state with *r* = 100 and *z* ≃ 2430.

**Figure 4.**
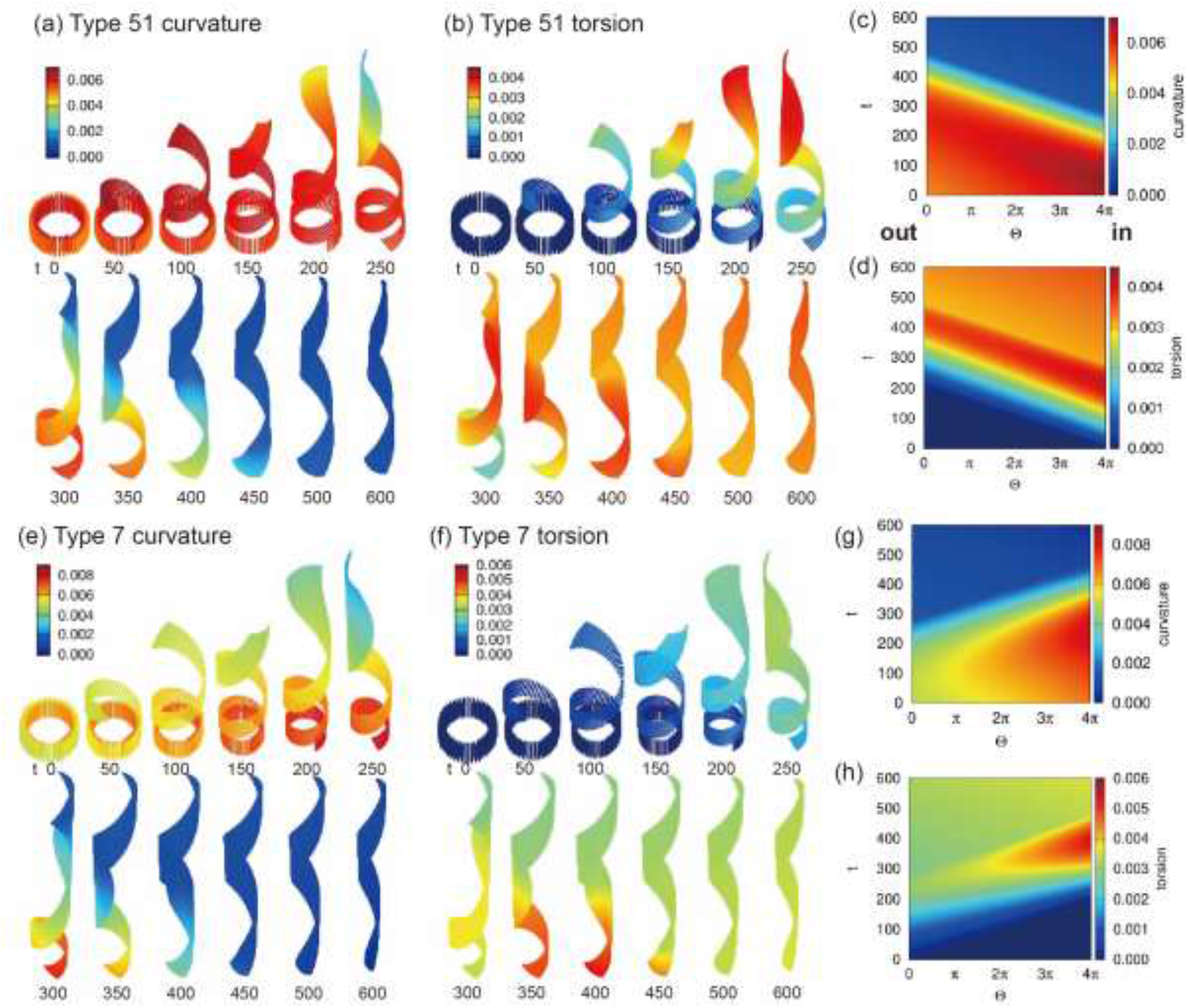
Macroscopic changes in curvature and torsion. (a, b) Spatio-temporal dynamics of curvature (a) and torsion (b) of type 51. (c, d) Curvature (c) and torsion (d) as a function of initially labelled coordinate *Θ* along the longitudinal direction and time t of type 51. (e, f) Spatio-temporal dynamics of curvature (e) and torsion (f) of type 7. (c, d) Curvature (c) and torsion (d) as a function of initially labelled coordinate *Θ* along the longitudinal direction and time t of type 7.

In summary, the deformation of the dynamically deforming end of the R-body is governed by a coupled process in which the curvature decreases through straightening and torsion is simultaneously enhanced via twisting.

### 2.3. Microscopic structural inference of lattice interval and lattice angle for type 51

We emphasise the significance of the DGT model for determining the microscopic structures of supramolecules. Using the initial set of type 51 lattice intervals as a representative example (figure 5*a*), we evaluated the deformed lattice intervals by applying the DGT model (figure 5*b*). We noticed that deformed longitudinal interval *δl*_in_ becomes directed such that it closely approaches the original height direction and deformed height interval *δd*_in_ becomes directed close to the original circumferential direction (figure 5*a,b*). As the initial rolled state is curved in the circumferential direction, outer lattice interval *δL*_out_ becomes larger as *δL*_in_ and thickness *δT* increases (figure 5*c*). We then confirmed that the DGT model controls *δl*_in_, *δd*_in_, and *δt* as a function of *g, m*, and *k*, respectively (figure 5*d,e,f*). Finally, we defined initial lattice angle *ϕ*, which is the inclined lattice interval of *δD*_in_ rotated by *ϕ* (figure 5*g*). This perturbation enables us to determine the extent to which the initial lattice angle affects deformed lattice angle *ϕ* (figure 5*h*). As a result, the resulting lattice angle *ϕ* can be completely evaluated as a function of *α* and *ϕ* (figure 5*i*). These results quantitatively determine the local lattice structure of the R-body.

**Figure 5.**
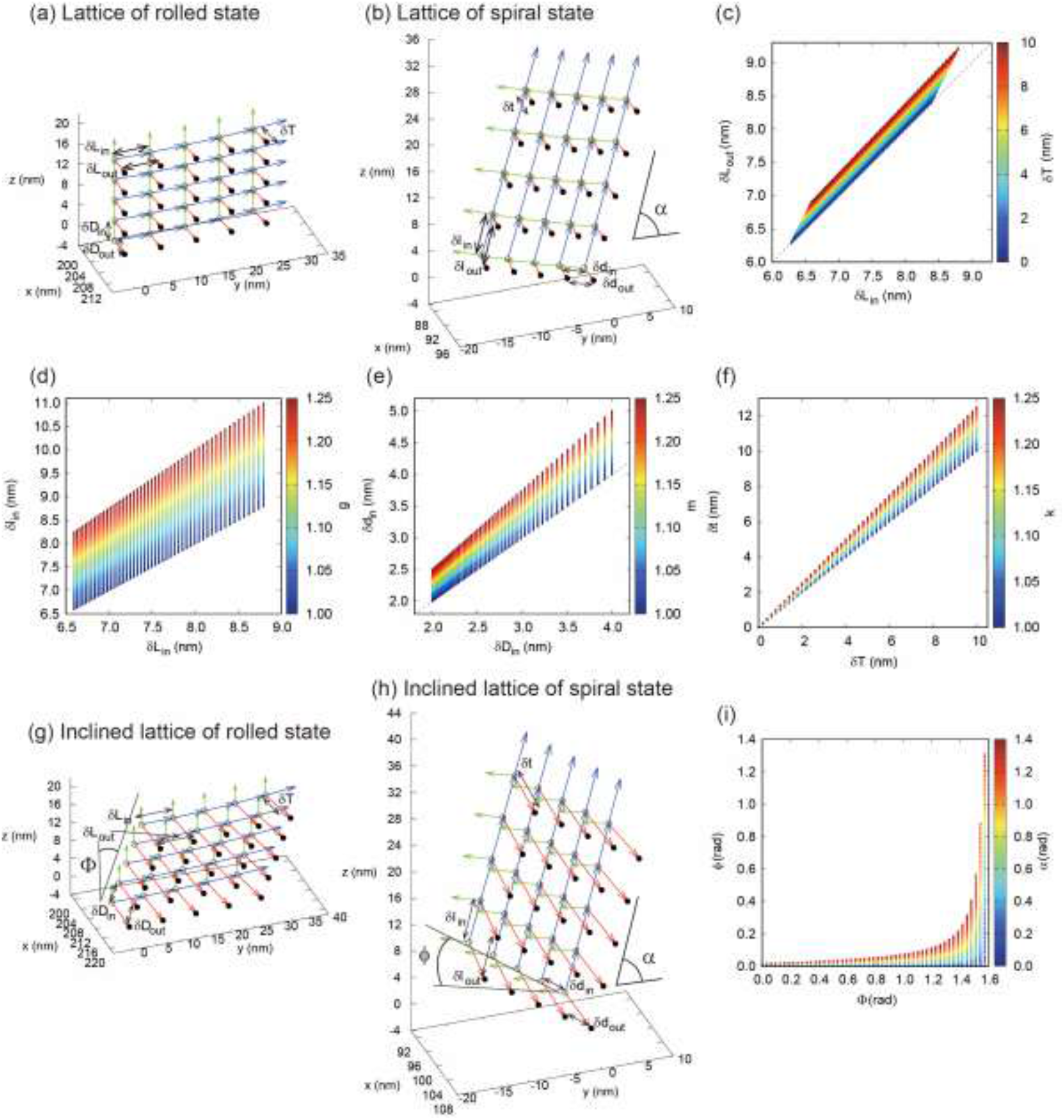
Microscopic change in lattice intervals of type 51. (a) Lattice of the rolled state. The longitudinal interval, height interval, and thickness interval are represented by *δL, δD*, and *δT* with subscripts in and out representing the inner and outer lattice, respectively. (b) Lattice of the spiral state. The longitudinal interval, height interval, and thickness interval are represented by *δl, δd*, and *δt* with subscripts in and out representing the inner and outer lattice, respectively. The inclination angle is *α*. (c) Outer length *δL*_out_ can be modified as a function of *δL*_in_ and thickness *δT*. (d) Deformed interval *δl*_in_ is transformed from *δL*_in_ according to the growth rate of length *g*. (e) Deformed interval *δd*_in_ is transformed from *δD*_in_ according to the growth rate of height *m*. (f) Deformed interval *δt* is transformed from *δT* according to the growth rate of thickness *k*. (g) Inclined lattice of the rolled state and angle *ϕ*. (h) Inclined lattice of the spiral state and angle *ϕ*. (i) Deformed lattice angle *ϕ* can be exactly determined by initial lattice angle *ϕ* and inclination angle *α*.

## 3. Discussion

In this study, we developed the DGT model to identify model parameters that express the dynamics of different types of R-bodies. The shape and deformation can be determined from the macroscopic (curvature and torsion) and microscopic (lattice interval and angle) morphological characteristics. These results provide a basis for the quantitative determination of the morphological structure of the R-body by establishing a bidirectional map between shape and deformation.

The significance of this study lies in presenting, for the first time, a quantitative theoretical framework for describing different types of reversible roll–spiral transformations of an R-body in a unified model. By employing the DGT, the spatio-temporal evolution of the curvature and torsion can be explicitly quantified, thus enabling a geometrical understanding of the deformation processes that were previously inferred mainly from morphological observations (Pond 1989). Importantly, the formulation of the DGT is the same for types 51, 7, and Pa, whereas the initial morphological setting precisely explains the different unrolling and tapered morphologies. These results demonstrate that types 51, 7, and Pa R-bodies share a common unrolling mechanism defined by the DGT, with a slight modification in the initial condition and onset of deformation. This implies the possibility that the local molecular deformation is almost the same for R-bodies of types 51, 7, and Pa.

Quantitative analysis showed that the curvature and torsion evolved in opposite directions during the roll–spiral transformation (figure 4), indicating that the principal curvature directions of the sheet undergo reorientation during deformation. Such geometric switching cannot be reliably inferred from the morphological appearance alone. This change in curvature is a phenomenon that resembles the mechanical buckling of the surface structure in leaves (Liang 2009); however, as the deformation progresses from the edge in contact with the liquid, it seems unlikely to buckle in the R-body, where the curvature and torsion evolve as intrinsic geometric variables encoded in the supramolecular lattice. Furthermore, by linking the changes in the macroscopic shape with the microscopic lattice structures, the framework allows molecular arrangements and structural transitions to be inferred. Notably, our results reveal that the subtle rearrangement of the surface grid can induce curvature switching, which in turn leads to roll–spiral macroscopic deformations guided by an important parameter, *e*. This hierarchical coupling demonstrates how local geometric reorganisation at the lattice scale can propagate and amplify across the assembly to generate global morphological transformations. Such multiscale amplification provides fundamental insights into the functional mechanisms of R-bodies and establishes a basis for future applications in the design of stimulus-responsive protein-based materials in soft matter physics (Hafner 2023).

It is worth noting that a methodological parallel with the iterative progress has been achieved in bacteriophage T4 studies. Characteristic helical contractions of the T4 tail sheath were first identified through macroscopic observations before high-resolution molecular structures became available (Amos 1975, Olson 1982). In response to these observations, geometric and continuum models have been developed to describe large-scale deformations and infer their three-dimensional organisation. These theoretical descriptions have shaped hypotheses regarding lattice arrangement and deformation mechanisms at the molecular level (Celotto 2003, Falk 2006). The continued accumulation of molecular and structural data (Kostyuchenko 2005) allows these hypotheses to be tested, refined, and integrated into a coherent understanding of the system (Aksyuk 2009, Fraser 2021). The resulting feedback loop between mathematical modelling and molecular structural analysis proved essential for constraining and validating the molecular structural models of the T4 tail sheath. In the case of R-bodies, extensive observations of the roll–spiral transformation exist (Pond 1989), yet a comparable quantitative framework is lacking. The proposed DGT model serves as a guideline for the deformation geometry of molecular organisation, thereby opening a similar pathway toward an integrative structural understanding of R-bodies.

This study involved the demonstration of a novel methodology for predicting the geometric constraints of chemical molecules using continuum mechanics. For example, the curvature and torsion could be extracted from the macroscopic morphological data and then used to derive the DGT. Alternatively, experiments could be performed to measure microscopic lattice deformation data and derive the DGT in an independent manner. Further meaningful applications of the DGT coupled with data lie in its ability to instantaneously compute changes in line with element lengths via the Cauchy‒Green deformation tensor, area changes using Nanson’s formula, and volume changes from the determinant (Goriely 2017), thereby providing concrete geometric constraints to link geometric measures to design principles and experimental design. The concept of correspondence between the shape and deformation is crucial, as it enables us to infer deformational dynamics by constructing continuum models from macroscopic or microscopic morphological changes on surfaces.

## Data accessibility

The data and computational code that support the findings of this study are available from the corresponding authors, S. Tsugawa and T. Ueno, upon reasonable request.

